# Beta-band dynamics during a naturalistic motor task: an OPM-MEG study

**DOI:** 10.64898/2026.07.27.740561

**Authors:** Joseph Gibson, Jessikah Fildes, Alan Kirby, Niall Holmes, Lukas Rier, Karen J. Mullinger, Matias Ison, Peter G. Morris, Elena Boto, Ryan M. Hill, Matthew J. Brookes

## Abstract

Beta oscillations are a fundamental feature of brain activity, linked to long-range connectivity within canonical networks, inhibition of sensorimotor cortices and the maintenance of a stable sensorimotor state in situations where the external world is predictable. The importance of beta oscillations in brain function is underscored by observations of their perturbation in neurological and psychiatric disorders. However, the precise role played by beta activity, particularly in mediating complex or skilful movements, remains incompletely understood. Here, we used a newly developed wearable optically pumped magnetometer-based magnetoencephalography (OPM-MEG) system to measure beta dynamics as participants learned to play a musical instrument. Twenty-two novice players took part in a study in which OPM-MEG data were recorded during two scanning sessions, while participants attempted to play a tune on a violin. Between the two sessions, participants received a violin lesson from an expert teacher. Results showed that robust data could be acquired during this naturalistic task, with beta oscillations decreasing in amplitude during movement and increasing upon movement cessation, as expected. Moreover, beta power whilst playing, in the motor and pre-motor areas, was significantly elevated after the lesson compared to before, and the movement-related modulation of beta amplitude was more pronounced after the lesson. These findings align with predictive coding models which suggest that beta amplitude should increase when individuals have greater certainty over the movements they carry out. Our study adds to an expanding literature on the role of beta oscillations and provides further evidence for the utility of OPM-MEG in naturalistic neuroscience.

## INTRODUCTION

Electrophysiological activity in the brain is dominated by rhythmic fluctuations across a wide frequency range (Pfurtscheller & Lopes da Silva, 1999). These oscillations play a key role in healthy brain function by coordinating neural assemblies. Specifically, brain regions communicate when their neural oscillations are aligned in phase; this creates time windows of high excitability that synchronize information exchange (Fries, 2005). Such ‘communication by coherence’ works on multiple scales, from local signalling (e.g. within a cortical column (Fries et al., 2001)) to long range coordination (e.g. across distributed networks (Hipp et al., 2012)). Neural oscillations in the beta (13-30 Hz) band are one of the most widely observed phenomena (Barone & Rossiter, 2021); they are strongest over sensorimotor regions, modulated by sensory or motor tasks, and have been implicated in connectivity within the sensorimotor network (e.g. (Brookes et al., 2011b; Hipp et al., 2012; Engel et al., 2013). The importance of understanding beta oscillations is underscored by their disruption in several neurological and psychiatric conditions; including Parkinson’s disease (Gross et al., 2001), multiple sclerosis (Barratt et al., 2017), schizophrenia (Robson et al., 2016) and autism (Buard et al., 2018). However, the precise functional role of beta oscillations remains unknown. Further, what is known comes mostly from laboratory-based studies using simple experimental protocols –e.g. finger movements. Developing a complete understanding of how beta oscillations, and the networks that they mediate, facilitate movement requires study of more complex paradigms, yet this is challenging with existing technologies. Here, we aim to demonstrate that new technology enables investigation of beta dynamics during a complex naturalistic motor learning task, in which participants play a musical instrument. The study of beta oscillations, particularly during naturalistic tasks, is challenging. Their electrophysiological origins mean they cannot be measured by techniques like functional magnetic resonance imaging (fMRI), which is limited to metabolic assessments of brain function. Electroencephalography (EEG) – the most commonly available functional neuroimaging modality – can measure electrophysiological function. However, EEG signals (which comprise electrical potentials measured at the scalp, that fluctuate due to changes in neural activity) are distorted spatially and diminished in amplitude by the high electrical resistance of the skull, limiting sensitivity and spatial resolution (Baillet, 2017). The EEG signal is also obfuscated by electrical interference from non-neuronal sources, especially muscles (Whitham et al., 2007; Muthukumaraswamy, 2013). This complicates EEG measurements carried out during naturalistic tasks, which often require movement. Magnetoencephalography (MEG) measures the magnetic fields generated at the scalp by electrophysiological activity in the brain. Since magnetic fields pass relatively undistorted through the skull, MEG offers better spatial resolution and sensitivity than EEG. MEG is also less sensitive to artefacts from muscles (Muthukumaraswamy, 2013; Boto et al., 2019). However, conventional MEG scanners rely on cryogenic magnetic field sensors, which must be fixed in rigid helmets and cooled with liquid helium. This means that participants must remain still during recordings. Consequently, whilst it is possible to perform simple motor tasks (e.g. finger movement) in a MEG scanner, more complex movements are difficult, or even impossible, in the restricted scanner environment.

In recent years, optically pumped magnetometers (OPMs) have emerged as a new way to measure MEG data (Brookes et al., 2022; Schofield et al., 2023). Unlike sensors used for conventional MEG, OPMs are small (12.4×16.6×24.4 mm), lightweight (∼4 g each), and do not require cryogenic temperatures. OPMs therefore get closer to the scalp, meaning they detect a larger signal (as signal amplitude decays with the square of distance from the brain) (Boto et al., 2016; Boto et al., 2017; Feys et al., 2022; Hill et al., 2024). Additionally, unlike cryogenic sensors which are in fixed arrays, OPMs can be mounted flexibly on the head, adapting for head size. This offers advantages for scanning people with small heads (e.g. children) (Hill et al., 2019; Rier et al., 2024; Rhodes et al., 2025). Most intriguingly (in the context of this paper) OPMs can be mounted into wearable helmets, meaning the sensor array moves with the subject. This (coupled with close control of background field) enables movement during a scan (Boto et al., 2018; Holmes et al., 2018). It therefore becomes possible to scan individuals who find conventional scanner environments challenging. Moreover, it becomes possible to collect MEG data as participants carry out naturalistic tasks (involving movement). There are already several studies exploiting this, ranging from seated head movements (e.g. (Rea et al., 2022)) to changes in posture (sitting/standing) (Sanders et al., 2025) and more expansive movement such as walking (N. Holmes et al., 2023; O’Neill et al., 2025). Because OPM-MEG offers the advantages of MEG (high resolution; lower sensitivity to artefact from muscle) in a wearable form factor (similar to EEG), this ostensibly provides the best platform for the investigation of beta dynamics during naturalistic motor learning tasks.

Here, we aim to show that a recently developed OPM-MEG device (Schofield et al., 2024) can be used to investigate beta dynamics as a participant learns to play a musical instrument. Specifically, we collected data in participants whilst they learnt to play a violin, both before and after a lesson with an expert teacher. Our primary aim was to show that, despite the obvious challenges of naturalistic movement whilst playing, high-quality MEG data can be captured. Our second aim was to investigate the effect of the lesson on beta dynamics. In general terms, beta oscillations are thought to represent the maintenance of stable sensorimotor predictions; beta amplitude tends to increase when the sensorimotor system is confident in its prediction of the outcome of a motor action, and decrease when updating of the prediction is required (Engel & Fries, 2010). There are several studies to support this. For example, in a task involving moving a joystick-controlled cursor to visual targets, Tan et al. (Tan et al., 2016) showed lower beta modulation in trials with large execution errors. Pollok et al. (Pollok et al., 2014) used a serial reaction time task to show that beta amplitude during movement was higher for learned sequences compared to random sequences. Separate work (Moisello et al., 2015) showed that prolonged practice of a motor task elevated beta oscillations. Predictive-coding theories interpret this as beta reflecting the precision of top-down predictions (Adams et al., 2013; Bastos et al., 2012). These previous studies, whilst not based on a naturalistic learning paradigm, enable generation of a hypothesis for the present study. We hypothesised that, after a lesson, we would observe an elevation in beta amplitude in the motor network whilst playing the violin, compared to before the lesson – reflecting an improvement in prediction error. Similarly, we expected that the modulation of beta amplitude when playing the violin (compared to rest) would be larger after the lesson compared to before.

## METHODS

### System design

The OPM-MEG system was constructed using 64 triaxial OPMs (3^rd^ generation; QuSpin Inc. Colorado, USA), each with equal sensitivity to magnetic fields in three orientations (Boto et al., 2022). The full array therefore allows 192 independent measurements of the MEG signalacross the scalp (i.e. 192 channels). Each OPM contained a ^87^Rb cell; a VCSEL laser; a photodiode, and three orthogonal sets of electromagnetic coils. The operational principles of these sensors is well established (for a review see e.g. (Schofield et al., 2023)) and will not be repeated in detail here. Briefly, laser light is projected through a ^87^Rb vapour inside the ce **l** and onto the photodiode. If the sensor is operated close to zero magnetic field, absorption of photons by the ^87^Rb atoms (measured as a drop in signal at the photodiode) becomes a sensitive marker of local magnetic field fluctuations. The OPMs were mounted in 3D-printed helmets (Cerca Magnetics Limited, Nottingham, UK) which allow an even distribution of sensors across the scalp (see Figure 1A).

**Figure 1:**
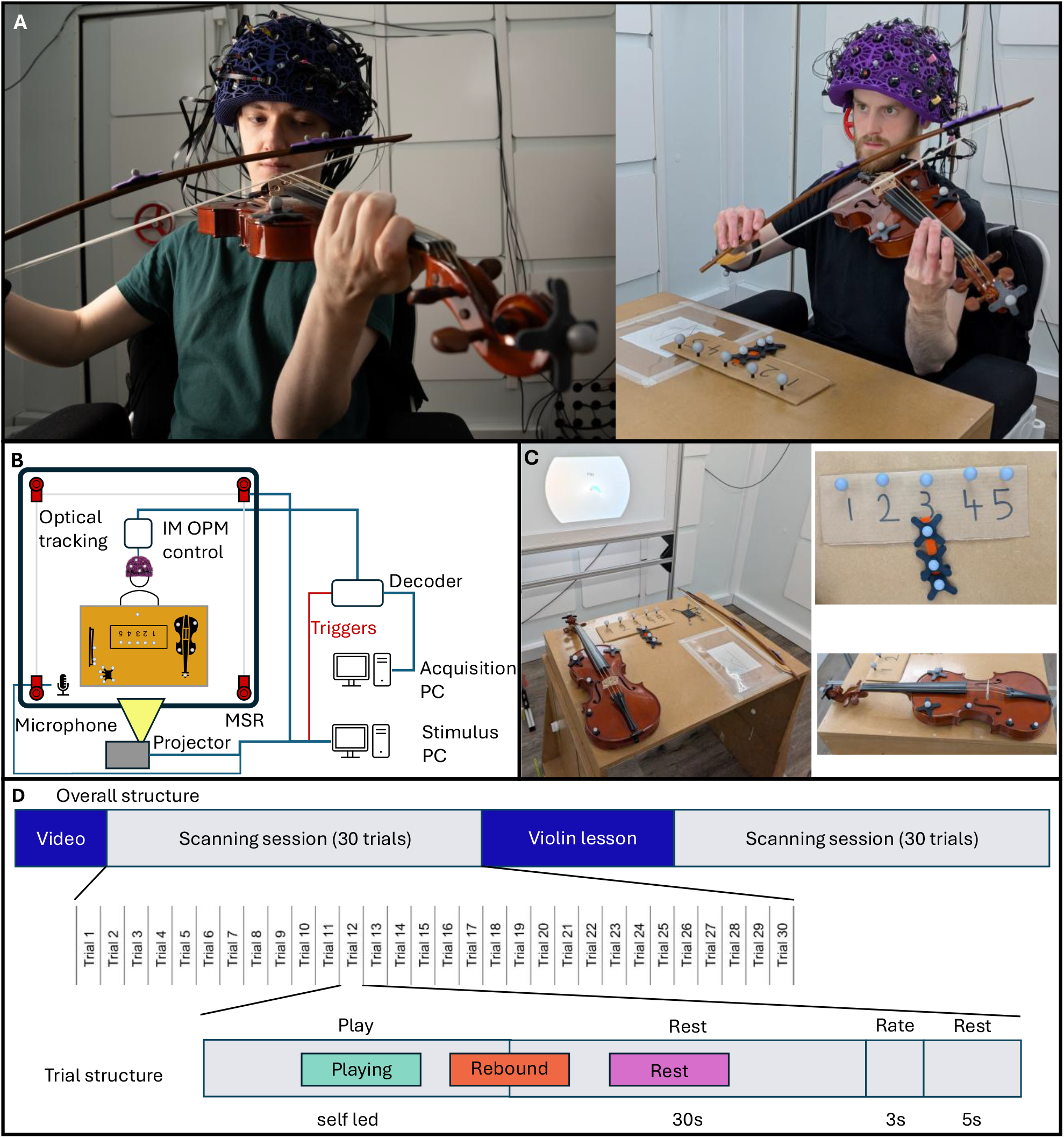
Experimental set up: A) Photographs of two participants undertaking the task. (Photos shown with written permission of the participants.) B) schematic drawing of the OPM-MEG system. C) Photographs showing detailed views of the equipment used in the experiment, including motion tracking markers, the modified violin and the rating scale. D) Schematic representation of the violin-playing task. The upper panel gives the overall structure of the participants visit whilst the lower panel gives the trial structure.

The array requires control signals to be sent, and output signals received, independently yet synchronously, to and from each of the 64 OPMs. This was enabled via an integrated miniaturised electronics unit (*“NEURO-1”*, QuSpin, Colorado, USA) (Schofield et al., 2024). The OPMs were operated in a “closed loop” mode whereby the measured field was fed back to the coils inside the sensor; currents through those coils were then updated in real time such that any change in externally generated magnetic field was offset by the coils (leaving the field at the cell, zero). The sensor output then effectively becomes the negative of the field produced by the coils. Without closed-loop operation, OPMs are limited to a dynamic range of ∼±1.5 nT (beyond which their output becomes non-linear). With closed loop, this is extended (e.g. to ±8 nT (Schofield et al., 2024)). The electronics unit was 0.36 x 0.2 x 0.06 m^3^ in volume, weighed 0.81 kg, and was wearable as a backpack (though for the present study it was simply attached to the back of a chair that the participants sat on). Each OPM is connected to the electronics unit via a lightweight cable, and the electronics unit itself has only two connections to the ‘outside world’ – a power cable and an Ethernet connection – the latter sends the digital output from all sensors to a decoder which integrates MEG signals with information on, for example, stimulus timing. The decoder is then connected to a PC, enabling data visualisation and storage.

The system was housed inside a 3 m x 3 m x 2.4 m magnetically shielded room (MSR) (MuRoom, Magnetic Shields Limited, Kent, UK). The room uses 4 layers of mu-metal shielding which attenuate DC and low frequency fields, and one layer of copper which attenuates high frequency fields. The room was equipped with degaussing coils which demagnetise its walls. This process (which takes approximately 60 s) reduces the remnant static (DC) field inside the room to ∼3-5 nT (Altarev et al., 2015). The room was also equipped with a microphone (to record audio) and a motion tracking system (OptiTrack, Flex13, Natural Point Inc., Oregon, USA) which uses an array of eight cameras to track the position of infrared retroreflective markers, which can be attached either to the participant, or to peripheral equipment. A single PC controlled the OPM array, data collection and degaussing. A separate PC controlled all stimuli delivered to the participants (see below), audio recording and motion tracking. A full system schematic is given in Figure 1B.

### Experimental Paradigms

Twenty-two participants (10 Male; 12 Female), with no previous experience of playing a violin, took part in the study. All participants provided written informed consent, and the study was approved by the University of Nottingham Faculty of Medicine and Health Sciences Research Ethics Committee. Out of the 22 participants, 16 self-reported as being right-handed; 6 self-reported as being left-handed, however all the participants were taught to play the violin right handed, for consistency.

The experimental protocol began with participants watching a video (2 mins 14 s long), in which an expert violinist was shown playing the first 4 bars of the tune “Twinkle Twinkle Little Star”. Participants then undertook the first of two scanning sessions, during which they attempted to play the same tune on a violin whilst MEG data were acquired. After this, participants attended a violin lesson, taught by the same expert featured in the video. Participants then underwent the second scanning session (identical to the first) in which they again attempted to play the tune (see also Figure 1D – upper panel).

Each scanning session comprised 30 trials. A single trial began with a visual stimulus instructing the participant to “play”. Following this prompt, the participant would pick up the bow and the violin (which was located on a table in front of them (see Figure 1C) and attempt to play the tune. Once they had finished, they placed the violin and bow back on the table. Replacing the instrument initiated the start of a rest period that lasted 30 s (during which the word “rest” was displayed on a screen). After the rest period, participants were asked to rate how well they had played the tune using a 5-point Likert scale; this feedback period lasted 8 s. (See also Figure 1D – lower panel.) Note that the trial length differed depending on how long it took subjects to play the tune. The average length of a scanning session was ∼35 minutes.

The lesson lasted approximately 30 minutes and comprised a standardised framework which included: 1) holding the violin; 2) holding the bow; 3) open string bowing; 4) correct finger spacing and pressure on the strings; and 5) playing the notes. These 5 steps were repeated to generate a reasonable outcome for each participant. The lesson ended with the subject playing the tune and the teacher simultaneously playing it, an octave higher, to allow the participant to gain a subjective measure of success.

The timing was such that the first scanning session, the lesson, and the second scanning session were carried out sequentially, on the same day. Precise timing varied from participant to participant (for practical reasons) but all those taking part completed the two sessions and lesson within ∼2.5 hours.

For all scanning sessions, visual stimuli were presented via projection (using a ViewSonic PX748-4K data projector) through a waveguide in the MSR onto a back projection screen, located ∼1 m from the subject.

The violin was specially adapted for the study: we removed the chin rest; replaced the steel core strings with gut strings; used a baroque bow (made entirely from wood and natural horsehair) and removed the adjusters (screw operated mechanisms attached to the tailpiece to fine tune the strings). These adaptations were to ensure that neither the violin nor bow contained metal parts, the movement of which could generate artefacts in MEG data. To accommodate the gut strings, the violin was retuned to baroque frequencies (e.g., the frequency of the ‘A’ string was reduced from ∼440 Hz to ∼415 Hz, and other strings altered accordingly). These changes had a minor detrimental effect on the quality of the instrument. However, the participants recruited for this study had not previously played a violin and so would be unlikely to recognise this. The tuning of the violin was checked prior to every scanning session.

### Data Collection

For each scanning session, the participant was seated on a patient support with a table (holding the violin and bow) positioned a comfortable distance in front of them. Once they were seated and had been given all relevant instructions the MSR door was closed, degaussing was initiated to demagnetise the inner metal walls of the MSR, and the OPM startup procedure was initiated. The degaussing and sensor startup took approximately 3 minutes. During this period, the participant was also shown the introductory video (in the first scanning session). Following this, the experiment was started.

We simultaneously captured data from the OPM-MEG system, the microphone (to record the sound of the violin) and the motion tracking cameras. For motion tracking, infrared retroreflective markers were attached to the violin, bow, and a ‘base station’ which was positioned on the table (see Figure 1C); all were modelled as rigid bodies. Motion tracking data from these bodies were fed, in real time, into the stimulus PC. The start of the rest period for each trial was triggered when the bow was replaced onto the table within a set distance of the base station.

Five ‘static’ retroreflective markers were also located on the table in front of the participant, next to cards containing the numbers 1-5 (see Figure 1C). During the feedback period of each trial, an additional ‘mobile’ retroreflective marker was moved by the participant between these numbers to indicate their self-assessment of how well they played the tune during each trial. The participant’s assessment (an integer score of 1-5) was determined by the proximity of the mobile marker to each of the 5 ‘static’ markers.

Retroreflective markers were also located on the OPM-MEG helmet, which was modelled as a rigid body and used to assess the extent of head movement during the experiment.

All motion tracking data were acquired at a sample rate of 120 Hz. All MEG data were acquired at a frequency of 375 Hz. The sound of the violin, recorded by the microphone, was fed through an audio interface (Focusrite Scarlett Solo) to the stimulus PC. These audio data were used to determine the moment at which the participant started to play in each trial. The motion tracking data, audio data and visual stimuli were synchronised to the MEG data by voltage pulses sent from the stimulus PC to all other systems.

Throughout the scanning session, the participant could contact the scanner operator via an intercom. Scanner operators could also see the participant on a video screen, via a camera located inside the MSR.

The scanning procedure was identical for sessions 1 and 2, with the only difference being that the video of the expert playing the tune was not shown in the second session.

### Coregistration to anatomy

Prior to their OPM-MEG scans, every participant had undergone an anatomical MRI (using an MPRAGE sequence implemented on either a Phillips 3 T Ingenia or a Phillips 7T Achieva MRI scanner). Following each OPM-MEG session, a structured light scanner(Einscan H, SHINING 3D, Hangzhou, China) was used to generate a 3-dimensional optical scan of the participants head with the OPM-MEG helmet in place. The subject’s face surface, extracted from the optical scan, was aligned to the equivalent surface extracted from the MRI. This resulted in a coordinate transform between the optical scan and the MRI reference frame, giving the helmet position relative to MRI. Combining this with the helmet sensor geometry (taken from the helmet 3D-printing process) gave information on the sensor locations/orientations relative to brain anatomy

### Data analysis

All data analyses were carried out using FieldTrip (Oostenveld et al., 2011) alongside bespoke algorithms written in MATLAB.

#### Preprocessing

Data were organised according to the Brain Imaging Data Structure (BIDS) for MEG (Niso et al., 2018). For all OPM-MEG data, an infinite impulse response notch filter was applied to remove powerline noise ( at 50 Hz, 100 Hz and 150 Hz). Data were then bandpass filtered into the 1-150 Hz band using a 4^th^-order Butterworth filter. Data were also processed using spatiotemporal Signal Space Separation (tSSS) (Taulu et al., 2005; Niall Holmes et al., 2023) to remove residual magnetic interference.

For each MEG channel, a power spectral amplitude plot was created (using Welch’s method (Welch, 1967)) and channels with a mean signal < 7 fT/sqrt(Hz) in the 60-80 Hz band (i.e. recording no signal) or a mean signal > 30 fT/sqrt(Hz) (i.e. a channel with high instrumental noise) were removed.

Trials containing high levels of interference werealso removed. This was achieved in two ways. First, the data from all channels, from each trial independently, were inspected visually and any trials containing high noise were removed. This allows identification and removal of trials with low-frequency interference but can miss interference at high-frequency. This is particularly relevant in naturalistic paradigms where movement is encouraged since magnetic artefact from muscles (e.g. in the shoulders or neck) typically manifests at high frequencies. To account for this, for each channel and each trial we derived a time-frequency spectrum (TFS) of the data. To do this, data from a single channel/trial were frequency-filtered into overlapping bands between 5 and 110 Hz (Specifically: 1.5-Hz wide bands, overlapping by 0.5 Hz, (i.e. 5- 6.5Hz; 6-7.5Hz; 7-8.5Hz Hz etc) up to 110 Hz). For each band, a Hilbert transform was applied to the filtered data to derive the analytic signal; the absolute value of the analytic signal was then calculated to generate a new timecourse showing the instantaneous amplitude of the band-limited signals – termed the Hilbert envelope, *A*(*t*, *f*). For all frequency bands, we also calculated a baseline oscillatory amplitude, *B*(*f*) as the mean (over time) of the envelope in the 0 s to 10 s time window (relative to the start of the rest period). We then computed the final TFS as *R*(*t*, *f*) = (*A*(*t*, *f*) − *B*(*f*))/*B*(*f*), which represents the fractional change in oscillatory amplitude for all frequencies relative to baseline. TFS’s were averaged across MEG channels, leaving a single TFS for each trial. These were inspected visually for every trial and, for any TFS showing high noise levels, the corresponding trial was removed. Example TFS’s resulting from this process are shown in Appendix 1. Following bad trial rejection, we also averaged the remaining TFS’s across trials, independently for each channel. Any channel showing high noise was also removed.

#### Source localisation and time frequency spectra

We applied beamforming (Van Veen et al., 1997 ; Robinson & Vrba, 1998) to generate images showing the spatial origins of task-induced beta modulation. We also used beamforming to derive a time-frequency representations of electrophysiological activity at locations of interest in the cortex (where locations of interest are identified from the peaks in the images) .

It is well known that periods of movementinduce a drop in beta amplitude (known as the movement-related beta decrease (MRBD)) whilst movement cessation causes an increase in beta amplitude (above baseline, known as the post movement beta ‘rebound’ (PMBR)). Beta amplitude only returns to baseline levels several seconds after the peak of the PMBR (Fry et al., 2016; Pakenham et al., 2020). Based on this, we defined three time windows of interest:

1. The *playing window* began at the point the subject played their first note (defined based on the audio recording made inside the MSR) and lasted for 10 s (during which the subject played the tune).
2. The *rebound window* began 5 s before the violin and bow were replaced (defined by motion tracking) and ended 5 s after they were replaced (i.e. was 10 s in duration).
3. The *rest window* began 10 s after the violin and bow were placed on the table following playing. This was also 10 s in duration.

Data were filtered to the beta (13-30 Hz) band (using a 4^th^-order Butterworth filter). Data covariance matrices were constructed independently for each time window and trial. These were averaged across trials leaving a single (beta-band) covariance matrix for each of the three time windows. These matrices were regularised using the Tikhonov technique with a regularisation parameter equal to 0.1% of the maximum singular value of the unregularized matrix. (Note that this relatively low level of regularisation was used to maximise the interference rejection properties of the beamformer (Brookes et al., 2008); this was considered advantageous in an experiment likely to cause significant interference – e.g. from muscle artefact.)

The brain was divided into cubic voxels with 4-mm isotropic resolution; for each voxel we generated a forward field based on the coregistration data, a dipole model of current in the brain, and a single-she **l** volume conductor model (Nolte, 2003). Beamformer weights, describing the optimal linear combination of MEG channels to represent beta-band activity at each voxel were then constructed using the forward field alongside the data covariance matrices (averaged over all three windows). For each voxel, we contrasted beamformer projected beta power in the playing and rest windows, using a pseudo-T-statistical approach (which computes the difference in power between the windows, normalised by twice the projected power in the rest window). For every voxel, source orientation was taken as the direction of maximum projected power (Sekihara et al., 2004). Plotting pseudo-T-statistics at each voxel generated functional images showing the spatial signature of beta modulation across the brain. These were averaged across participants.

For the locations showing the largest task-induced beta modulation (taken from pseudo-T-statistical images, averaged over subjects and sessions) we undertook time-frequency analyses. Beamformer weights were rederived using covariance calculated in the 1-150 Hz band, again averaged over all three windows and regularised as above (The wider band allowed us to construct TFS’s over more than just the beta range). These weights were then used to derive a broadband estimate of electrophysiological activity at the locations of interest (termed a virtual electrode (VE)). For each location, we computed a trial-averaged TFS (based on the Hilbert envelope – see above) for each of the three time windows. For all three windows, the TFS data were normalised relative to the mean oscillatory amplitude in the rest window. TFS data were computed independently for locations in left and right motor cortex and for data recorded in each session (i.e. before and after the lesson). For visualisation, the data were averaged across subjects.

We hypothesised that beta modulation would differ before and after the lesson. This was based on the idea that participants should experience greater certainty about their movements after the lesson compared to before. To test this statistically, for each subject we took the average beta amplitude during the playing window (relative to rest) for the session before the lesson and subtracted this from the same metric computed after the lesson. This was repeated for all participants and we used a Wilcoxon sign-rank test to determine if the resulting set of values were significantly different from zero. This test was carried out independently for locations in the left and right hemisphere, and the threshold for significance was recalculated to account for this (i.e. Bonferroni correction).

#### Relative Spectral Power Analysis

The above analyses only account for beta-band *modulation* with the task (i.e. changes in amplitude relative to the rest window). We also wanted to investigate changes in absolute spectral power between scanning sessions and, to this end, we undertook a beamformer-based power analysis.

The brain was parcellated into 82 cortical regions, defined by the MarsAtlas (Auzias et al., 2016). This was done by first defining the MarsAtlas in the space of the MNI template (ICBM152) brain. This template was then warped to the MRI of every participant using FLIRT in FSL (Jenkinson et al., 2002; Jenkinson & Smith, 2001). The same transform was applied to the MarsAtlas to define the 82 regions in each individual’s anatomical space. The coordinates of the centre of mass (centroid) of each region were then determined. OPM-MEG data were filtered into the beta-band. For each region centroid, we used data covariance matrices to derive beamformer weights, independently for each time window (using 1-150 Hz data, with regularisation as above) (Barratt et al., 2018). The weights were then used to reconstruct beta-band VE signals, again independently for each time window. For each VE constructed, we used Welch’s approach to derive the spectral content of the signal and integrated across frequency to derive the total spectral power in the beta-band. These calculations were then repeated, identically, but using broadband (rather than beta-band) filtered data. We then computed the ratio of beta power to broadband spectral power, providing a final metric of *relative beta power* (Rier et al., 2023). Doing this for each region, and independently for our three time windows, enabled construction of images showing the distribution of relative beta power across the brain and how it changes during the task (i.e. across our three time windows).

For each time window, we compared relative beta power before and after the lesson. To avoid a multiple comparison problem (across 82 regions) we assumed that beta change elicited by the lesson would be largest in the sensorimotor network. We therefore averaged relative beta power in the left and right dorsolateral and dorsomedial motor areas, the left and right dorsolateral and dorsomedial somatosensory areas and the left and right dorsolateral and dorsomedial premotor regions (i.e. averaging across 12 regions in total). For each participant, this left six metrics of *sensorimotor network relative beta power;* one for each time window before the lesson, and one for each time window after the lesson. We then subtracted the metrics before the lesson from those computed after the lesson to derive the difference in relative beta power between sessions. These values were compared statistically using a Wilcoxon sign-rank test and we corrected for multiple comparisons (across the three windows) using Bonferroni correction (i.e. the threshold for significance was altered to 0.05/3).

We also tested for a relationship between the change in relative beta power and participant performance. The latter was measured as the change in the self-assessment scores representing how we **l** each participant played the tune. Specifically, the self-assessment scores for all trials before the lesson, and all trials after the lesson, were averaged independently and the difference between the averages calculated. This resulted one difference score per participant. We then calculated the Pearson correlation (over participants) between this behavioural metric and the change in relative beta power, for each window, hypothesising that participants with a higher beta change would exhibit a larger improvement in self - assessment. The threshold for significance was adjusted to account for multiple comparisons.

#### Functional Connectivity Analysis

We used amplitude envelope correlation (AEC) (de Pasquale et al., 2010; Brookes et al., 2011a; Hipp et al., 2012) to characterise functional connectivity between all pairs of regions in the MarsAtlas. Briefly, beta-band filtered data were used to construct beamformer-derived VE signals for all MarsAtlas region centroids (as above). For each pair of brain regions, pairwise orthogonalisation was applied to reduce the effect of source leakage (Brookes et al., 2012; Hipp et al., 2012). Following this, the Hilbert envelopes for the two regions were calculated and the Pearson correlation between the envelopes calculated to quantify functional connectivity (i.e. if 2 regions envelopes are correlated they are said to be functionally connected). This was repeated for all region pairs, resulting in a whole-brain connectome matrix showing how each region is connected to every other region. Connectome matrices were constructed independently for each task trial and for each time window and results averaged across trials, leaving three matrices representing the playing, rebound and rest windows. Connectome matrices were averaged over participants for visualisation. In addition, we derived a threshold for the strongest 5% of connections for all 3 time windows. This was done for data recorded before and after the lesson, and matrices thresholded with the higher of the two values. The strongest connections were plotted as lines in a glass brain, for visualisation. We also calculated connectivity strength for each region (i.e., how connected a region is to every other region – calculated as the sum of all elements along a row (or column) of the matrices). This was also plotted on the glass brain.

We aimed to statistically compare functional connectivity in the sensorimotor network before and after the lesson. Here, we summed connectivity strength across the 12 regions in the sensorimotor network, providing three metrics of “sensorimotor network connectivity” (one for each time window), before and after the lesson. We then derived the difference in sensorimotor network connectivity between sessions. These values were compared (as above) using a Wilcoxon sign-rank test and corrected for multiple comparisons using Bonferroni correction. Again, we tested for correlation between changes in functional connectivity and the averaged change in participant’s delf assessment scoring, using Pearson correlation.

## RESULTS

Data were successfully recorded in 19 out of 22 participants. From the remaining 3 participants, the trigger channels failed in 2, making it impossible to process the data. In 1 participant, data were unusable due to high levels of noise. 19 participants were therefore used for the final analysis.

Our preprocessing steps removed 13 ± 8 channels and 7 ± 6 trials (mean and standard deviation across participants) in scanning sessions carried out before the violin lesson. We removed 13 ± 6 channels and 6 ± 5 trials in scanning sessions carried out after the lesson. Channel removal was mostly due to faulty cables in the system, whilst trial removal was due to excess interference caused by muscles.

Head movement was tracked throughout the scanning sessions using the motion tracking cameras. For each participant and scanning session, we quantified the maximum distance that the centre of the head moved from its mean location. This was found to be 41 ± 29 mm before the lesson (mean and standard deviation across participants) and 68 ± 50 mm after the lesson. It is noteworthy that this degree of head movement would not be possible in a conventional MEG system. We used the standard deviation of the distance from the centre of the head from its mean location over the whole session to quantify the level of motion present in a scan. This was found to be 0.015 ± 0.007 mm before the lesson and 0.014 ± 0.006 mm after the lesson. We found no measurable correlation between head movement carried out by a participant and the number of rejected trials (Pearson correlation 0.20, p = 0.23).

The self-assessed performance rating was 2.8 ± 1.4 before the lesson, and this increased to 3.1 ± 1.3 after the lesson. This suggests that the lesson had an effect on the ability of the participants to play the violin, however the improvement was not statistically significant according to a Wilcoxon sign-rank test.

### Source localisation and time frequency spectra

Figure 2A shows the participant-averaged pseudo-T-statistical images depicting the areas of largest beta-band modulation between the playing and rest time windows. The image before the lesson is shown on the left and the image after the lesson on the right. In both cases, the largest beta modulation with the task is measurable in the left and right primary sensorimotor regions and is negative; this indicates the expected drop in beta power in the sensorimotor regions whilst moving to play the instrument, compared to rest (i.e. the MRBD). The peak locations themselves are in similar regions for both scanning sessions, and in both sessions the degree of beta modulation was largest in left hemisphere compared to right (perhaps reflecting the cohort was mainly right handers). Interestingly however, the pseudo-T-statistics (which essentially measure the signal to noise ratio of the beta change due to movement) tended to be larger (i.e. more negative) after the lesson compared to before. (Specifically: In the left hemisphere the pseudo-T-statistics were -0.14 ± 0.07 (before the lesson) compared to -0.15 ± 0.07 (after the lesson); p = 0.0015 (Wilcoxon sign rank test). In the right hemisphere the pseudo-T-statistics were -0.13 ± 0.07 (before the lesson) compared to -0.14 ± 0.07 (after the lesson); p = 0.1).

**Figure 2:**
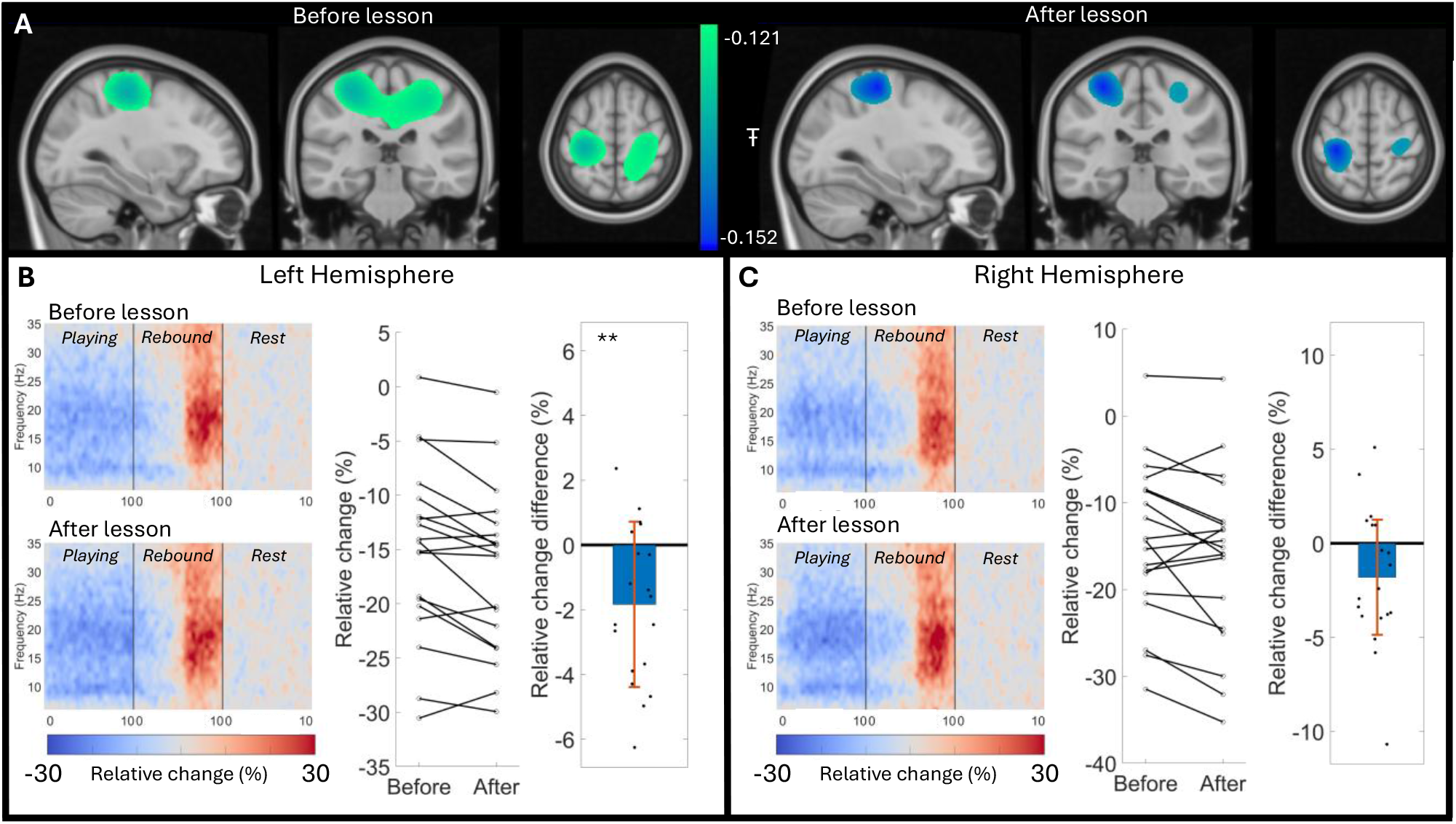
The spatiotemporal signature of movement related beta modulation whilst playing the violin. A) Participant-averaged pseudo-T-statistical images showing the locations of largest beta-band change when playing the instrument compared to rest. Left: before the violin lesson; Right: after the lesson. B) Left-hand panel: Time frequency spectrograms showing the time evolution of neural oscillations across the alpha, beta and low gamma frequency range, in left sensorimotor cortex. Upper plot shows before the lesson; lower plot shows after the lesson. All three time windows (playing rebound and rest) are normalised with respect to the mean oscillatory amplitude in the rest window , and are shown concatenated. Centre panel: mean movement related beta decrease in the playing window compared to rest, plotted for each subject before and after the lesson. Lines join the same subject in the two sessions. Right-hand panel: Differences in movement related beta decrease before and after the lesson. Statistical analysis shows the median of the distribution is negative (p = 0.007). C) Equivalent to (B) but for right sensorimotor cortex. Note here that a similar pattern is observed, however the differences before and after the lesson are no longer significant (p = 0.059).

In Figure 2B, the left-hand panel shows TFSs for the left primary sensorimotor region. Data for all three windows (playing, rebound and rest) are shown concatenated (normalised relative to rest). The upper panel shows data before the lesson and the lower panel shows data after the lesson. Blue indicates a loss in oscillatory amplitude (measured as a percentage relative to rest); red indicates an increase. The expected MRBD during playing is apparent, alongside an increase in beta amplitude above baseline (the PMBR) during the rebound window. This is the case for data recorded in both scanning sessions.

The centre panel of Figure 2B compares the amplitude of the MRBD before and after the lesson. Here, beta amplitude has been averaged within the playing window, and each data point represents the result from a single participant. The lines connect data for the same individual before and after the lesson. Notice that, on average, the lines slope down (i.e. the MRBD is more negative after the lesson). These data are further explored in the right hand panel of Figure 2B, which shows the differences in MRBD before and after the lesson. This is again measured as a percentage (i.e. in a subject where movement causes a 20% drop in beta amplitude before the lesson, and a 22% drop after the lesion, this difference would be -2%). In the Figure, individual data points represent each subject, whilst the mean and standard deviation are shown by the blue bar and red error bar respectively. Statistical analysis showed that the data likely come from a distribution with a median less than zero (p = 0.007) indicating that the MRBD was larger after the lesson compared to before. However, it should be emphasised that this effect is small compared to both the effect of movement on beta amplitude, and the inter-individual differences. Figure 2C shows equivalent data to Figure 2B, but for the right hemisphere; these data show a similar trend but the difference in MRBD before and after the lesson failed to reach statistical significance (p = 0.059).

### Relative Spectral Power Analysis

Whilst Figure 2 shows how beta amplitude modulates from rest throughout the task, Figure 3 extends this by including an assessment of underlying beta power (i.e. taking into account the baseline value). The brain images in the left and centre columns show the spatial distribution of relative beta power (i.e. the proportion of the overall spectrum that exists in the beta-band) across the MarsAtlas regions. The top, middle and bottom rows show the playing, rebound and rest windows respectively. The left-hand column shows the case before the lesson whilst the centre column shows data after the lesson. In the right-hand column, the datapoints show the average change in relative beta power in the sensorimotor network, between the two sessions, for each subject. The blue bar shows the mean value (across subjects) and the red error bar shows standard deviation. The 12 regions included in the sensorimotor network are shown by the inset image, with the different colours showing how much the relative beta power in each of the 12 regions changes between the two sessions (averaged across all participants – see also Appendix 2).

**Figure 3:**
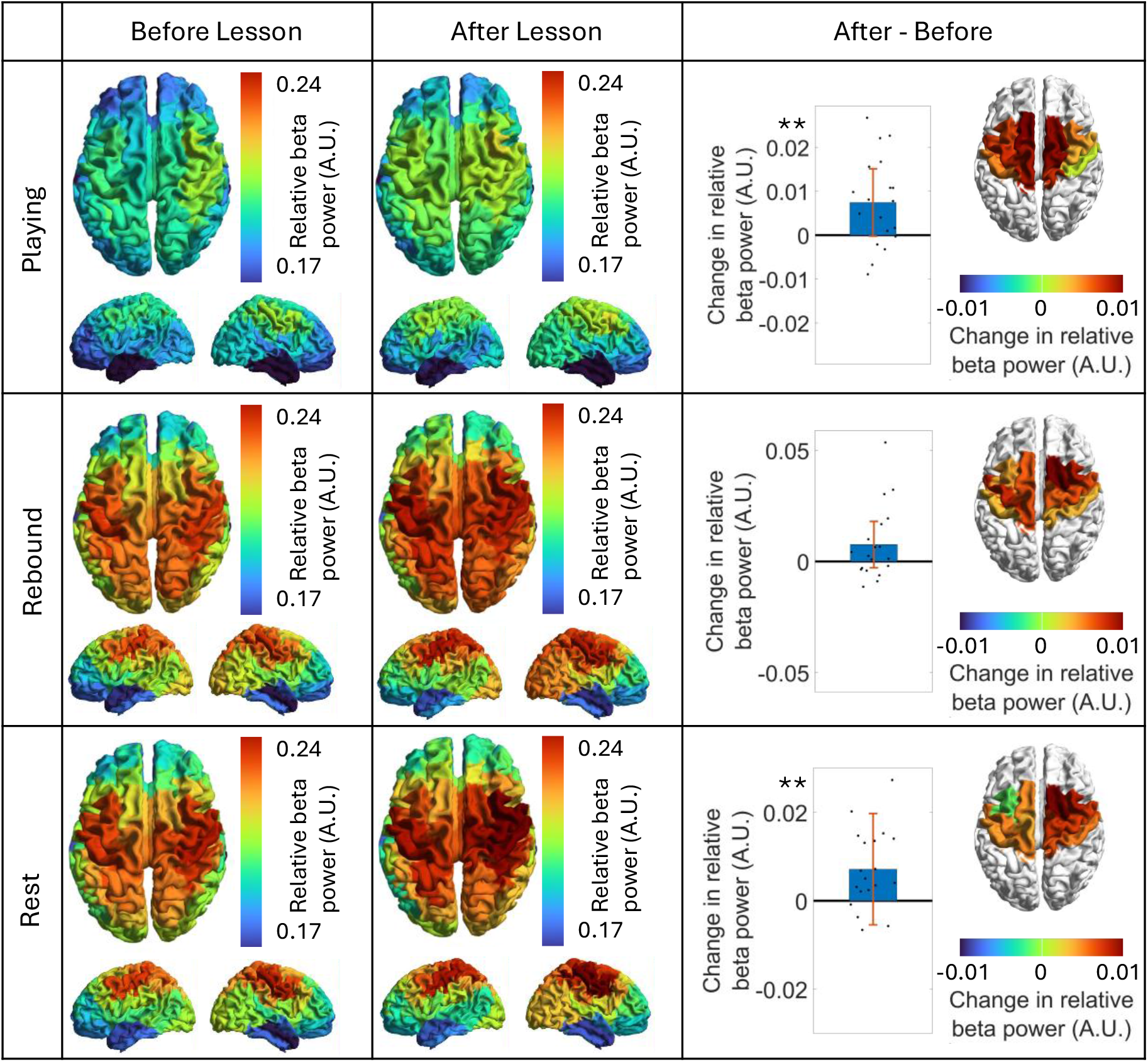
Beta-band power before and after the violin lesson. Left column: Images showing the spatial distribution of relative beta power before the lesson. Centre column: Images showing the spatial distribution of relative beta power after the lesson. Right column: The power difference between the two sessions, averaged across regions within the sensorimotor network. Data points show values for each participant; the bar shows the mean and error bar shows standard deviation. ** indicates a statistically significant difference between session (p < 0.05, corrected for multiple comparisons). In the inset image, the colours show the change in relative beta for each of the 12 regions in the sensorimotor network. The upper row shows the playing window, the centre row shows the rebound window and the lower row shows the rest window.

Looking at relative beta power across the three time windows, modulation of beta activity by the task can be seen clearly, with lower beta power during the playing window compared to rest. Note that, whilst the rebound and rest windows look similar, the rebound window actually contains the final few seconds of movement and so integrates a period of reduced beta and a period of enhanced beta.

More interestingly, whilst playing the violin (i.e the top row) the images appear to show a clearer motor network after the lesson compared to before. This is supported statistically; relative beta power averaged over the sensorimotor network regions increased by ∼0.0074; a change of ∼3.76% compared to beta power before the lesson. This increase was significant (p = 0.0079). During the rest window (bottom row) there was also a significant (p = 0.0055) increase in relative beta power after the lesson compared to before. The absolute increase was similar in magnitude (∼0.0071) but smaller as a percentage change (∼3.16%) between sessions (due to the larger overall beta power in the rest window. There was no measurable change in relative beta power during the rebound window.

Interestingly, a post hoc analysis (see again Appendix 2) showed that within the sensorimotor network, the individual MarsAtlas regions showing the largest change in beta tended to be the bilateral dorsomedial premotor areas during the playing window, and the right-lateralised dorsomedial and dorsolateral motor and premotor cortices during the rest window. The weakest effects were in left and right dorsomedial and dorsolateral somatosensory cortices.

During the playing window, Pearson correlation between the change in beta power across the sensorimotor network (before and after the lesson) and the change in the participants self -assessment of how well they played the tune (averaged across trials before and after the lesson) was 0.30. Although not significant using a two tailed test, (p = 0.071); based on a hypothesis that those subjects with a larger beta change will also have a larger change in self-assessed performance (i.e. where the direction of the change ostensibly means that one could carry out a one-tailed test) this would be significant (p = 0.036). A post hoc test, applied independently to the 12 brain regions in the sensorimotor network showed significant positive correlations (using a 2-tailed test) between beta power change and participant self-assessment in the left dorsolateral motor cortex (p = 0.04); the left dorsomedial somatosensory cortex (p = 0.034); the Left dorsomedial premotor area (p = 0.014) and the right dorsomedial premotor area (p = 0.02). Importantly, these findings would not survive a multiple comparison test but nevertheless show indication of a positive trend (data shown in Supplementary Figure S1).

We also measured Pearson correlation between change in beta power between sessions, and the difference in head movement between sessions. However, we found no significant correlation during either the playing (r = 0.07; p = 0.77) rebound (r = 0.11; p = 0.64) or rest (r = -0.21 p = 0.40) windows.

### Functional Connectivity Analysis

Results of our connectivity analyses are shown in Figure 4. In the left and centre columns, the connectome matrices show the strength of functional connectivity across all possible pairs of MarsAtlas regions (i.e. each matrix element shows the amplitude envelope correlation of beta signals between two different regions). A darker red indicates a stronger connection. In the ‘glass brain’ plots, the red lines show the strongest connections between regions either before or after the lesson, whilst the diameter of the blue dots show connectivity strength to each region (i.e. how connected that region is to all other regions in the brain). In the right hand column, the datapoints show the average change in connectivity strength in the sensorimotor network, elicited by the lesson. (Again, the regions included in the sensorimotor networks are shown inset.) The blue bar shows the mean and the error bar shows standard deviation.

**Figure 4:**
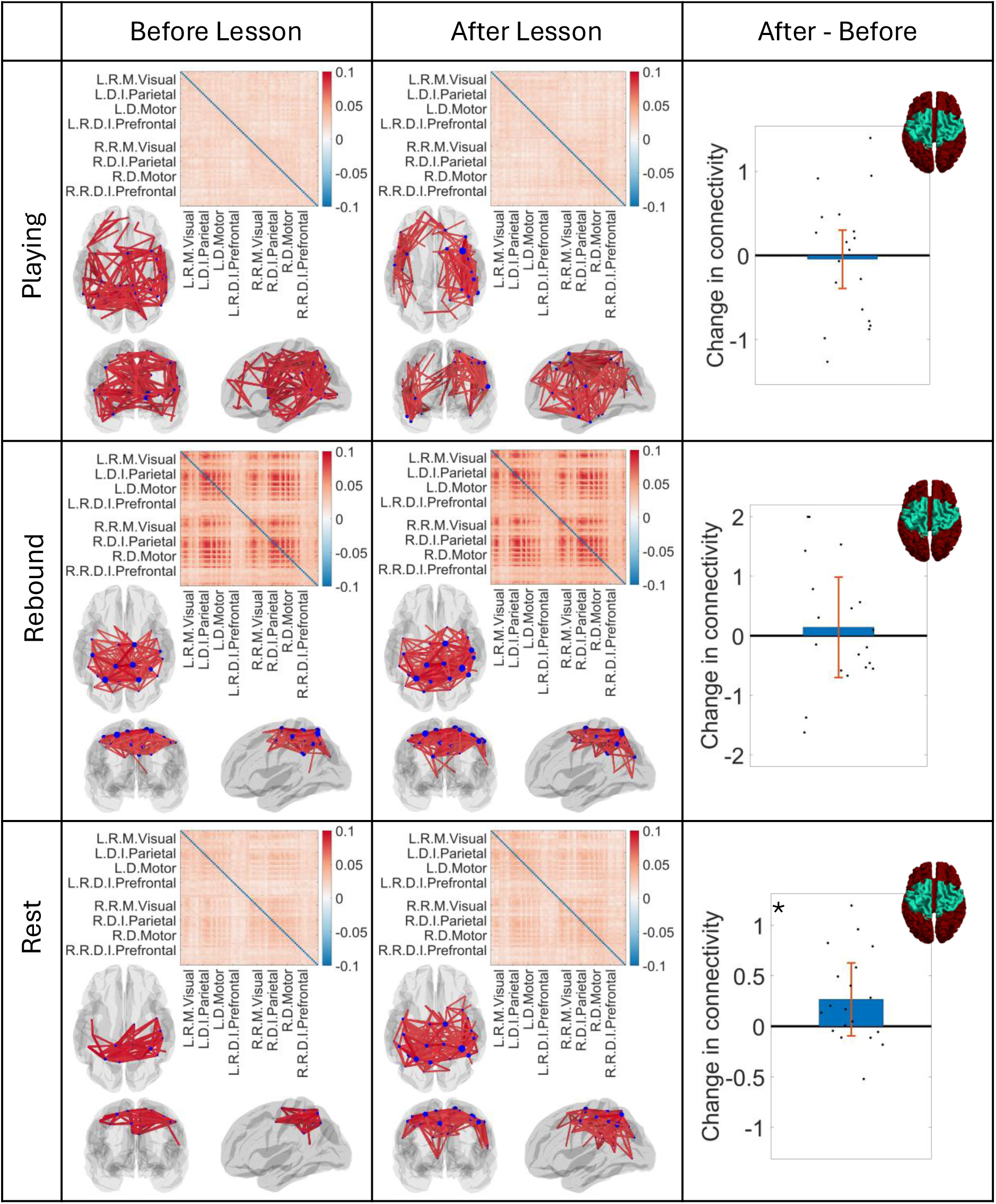
Beta-band functional connectivity before and after a violin lesson. The connectome matrices show functional connectivity between all possible pairs of MarsAtlas regions. The glass brain shows the spatial distribution of the strongest connections. The Left column shows the case before the lesson; The centre column shows the case after the lesson. The upper row shows the playing window, the middle row shows the rebound window, and the lower row shows the rest window. The right column shows the difference in connectivity strength in the sensorimotor regions (shown inset) between the two sessions. Data points show values for each subject; the bar shows the mean and error bar shows standard deviation. * indicates a trend (p < 0.05 but not corrected for multiple comparisons).

In the rest window, beta connectivity was found to be largest between regions that comprise the sensorimotor network and this was true of data acquired both before and after the lesson. During the rebound window, connectivity within this network was markedly stronger than during the rest window, but the spatial pattern remained similar. During playing, connectivity in this network breaks down and the strongest connections become more widely distributed across the brain (likely reflecting noise rather than the dominance of a new widespread network).

Both during the playing and the rebound window, there was no measurable change in functional connectivity between the two scanning sessions. During the rest window there was a trend for higher connectivity after the lesson compared to before. However, this trend (p = 0.024) failed to reach a revised threshold for significance following multiple comparison correction.

We found no measurable relationship between the connectivity change between sessions, and the participant self-assessment score changes.

## DISCUSSION

Beta oscillations in the sensorimotor system perform several putative roles. First, they impose inhibition on the primary sensory and motor cortices. The evidence for this is widespread. Beta oscillations drop during movement (the MRBD) and increase on movement cessation (the PMBR), both of which imply that high beta amplitude opposes movement. There is also more subtle evidence – for example switching sensory attention from the left to the right hand causes an increase in beta amplitude in right primary somatosensory cortex and a simultaneous decrease in left somatosensory cortex (Bauer et al., 2006; Rivero et al., 2025; Tanner et al., 2025). This is again consistent with increased beta amplitude reflecting inhibition. Second, beta oscillations carry a forward model relating to the consequences of movement – this is the so-called ‘predictive coding’ theory and again evidence for this exists. For example, there is typically stronger beta activity when movement is predictable (i.e. when the “status quo” is expected) and lower beta amplitude when a change is needed in predicting the outcome of a movement (Engel & Fries, 2010). Thirdly, beta oscillations are thought to help maintain long-range communication in the brain. This is evidenced by numerous studies (e.g. (de Pasquale et al., 2010; Brookes et al., 2011b; Hipp et al., 2012)) showing that canonical brain networks can be delineated from MEG data by assessing the spatiotemporal structure of beta-band signals. Recent years have also generated research showing conclusively that beta oscillations are not always smooth, continuous oscillations, but are in fact formed from punctate, transient events (Sherman et al., 2016; Little et al., 2019). This supports the idea that the inhibitory effect of beta “oscillations” can be transient. Our findings here are largely supportive of all these ideas.

Our primary findings were: 1) The overall beta power, both when playing the violin and in the rest periods between playing, was elevated after a violin lesson (when participants had been trained) compared to before (when they had little idea of how to play). 2) The change in beta amplitude elicited by playing the violin (i.e. the MRBD) was higher (more negative) after the lesson compared to before. Naively, these findings may seem at odds with each other – i.e. the beta decrease during playing (MRBD) was more negative, yet beta power during playing was higher. However, this can be explained because the MRBD is measured relative to the rest window, and the resting beta power was also significantly higher after the lesson. So, after the lesson, a larger beta modulation elicited by playing rides on top of a higher overall beta amplitude.

Elevated beta power following the lesson fits well with the predictive coding model, that beta is enhanced when there is greater certainty about the movements being undertaken, and furthermore it agrees with previous studies (Pollok et al., 2014; Moisello et al., 2015; Tan et al., 2016) which all show similar results. The finding also fits with the putative role of beta as an inhibitory process. For example, it is tempting to speculate that, whilst playing the violin, the movements required are facilitated by bursts of inhibition that stop unwanted motion. By extension, as a participant improves at the task , one would expect an increased beta signal during playing, and this is what we see elicited by the lesson. Beta amplitude was also increased between periods of playing and this could be accounted for if, for example, increased beta oscillations within the motor network helped reinforce the positive outcome of a task (again maintaining a ‘status quo’).

We observed some (limited) evidence for a correlation between beta power change and behaviour (the latter subjectively measured by the participants self-assessment of how well they played the tune). This finding was strongest during the playing window, and in the left and right dorsomedial premotor areas – areas known to be involved in the planning of movement. These findings did not survive multiple comparison correction and so are only reported as a trend. Nevertheless, they would fit with the idea of the predictive coding model for beta oscillations, where those individuals with greater beta modulation between the two sessions improved their playing the most.

The network aspect of beta oscillations was also observable in our study, with networks of functional connectivity clearly delineated by our connectivity analysis, and spatial modes centred on the sensorimotor regions. Interestingly, this network breaks down during the playing window (see Figure 4). Whilst there were no significant changes in functional connectivity after, compared to before the lesson, there was a trend for increased connectivity within the sensorimotor system after the lesson, which could be explained by the notion of reinforcement following a more precisely carried out task.

The primary aim of our study was to determine whether a recently developed 192-channel OPM-MEG system could be used to capture high-fidelity data on beta dynamics throughout a naturalistic task. As discussed in our introduction, measurement of beta oscillations is challenging: Techniques like fMRI or PET cannot be used, not only because electrophysiological signals are inaccessible to them, but also because space and movement constraints inside MRI and PET systems (and the high acoustic noise levels of MRI) would make playing a violin (and indeed many other naturalistic tasks) impossible. The task could be carried out during EEG acquisition, and EEG can be used to measure beta oscillations. However, EEG is extremely susceptible to high levels of interference from muscle activity, and it was felt that the large amounts of shoulder and neck movements required to play a violin would likely lead to significant artefacts in EEG, which may obscure signal of interest. In addition, spatial resolution in EEG is limited, consequently gaining sufficient resolution, for example to differentiate MarsAtlas regions for our power and functional connectivity analyses, is a significant challenge. Conventional MEG does offer high sensitivity to beta oscillations and good spatial resolution, but the static nature and large size of conventional MEG instrumentation would make carrying out the task very difficult. For this reason, we posit that OPM-MEG is the only technology that provides the optimum combination of performance and practicality that is required to undertake the current study.

There are several design considerations, specific to the OPM-MEG device used, which make it an ideal platform for naturalistic neuroscience. First, the system is constructed around a helmet that is small (total helmet thickness 32 mm), lightweight (∼900g), relatively unobtrusive, and accommodates natural movement during scanning. Participants were able to sit comfortably and play the violin without being hindered by the equipment. Second, the paradigm required significant head movement (up to ∼70 mm, measured via motion tracking). Even after degaussing, there is a remnant background magnetic field of ∼3-5 nT inside our MSR, and as OPMs move through this static field, they “see” a changing magnetic field which can be large (we measured it up to ∼ 5 nT). Such field shifts would be sufficient to take a standard Rb-based OPM outside its dynamic range (i.e. the range over which its output is a linear function of magnetic field). However, here we were able to exploit closed-loop measurements to ensure linear operation even with large field shifts (Schofield et al., 2024). Third, we employed triaxial OPMs (i.e. with MEG sensitivity along three axes). This has 3 distinct advantages over both single-axis sensors (as used in conventional MEG and some OPM-MEG systems (Rea et al., 2022)) and dual-axis devices. 1) It maximises channel density, ensuring the largest possible signal is captured by the array, which is known to improve results following source localisation (Hill et al., 2024). 2) It enables optimal separation of the signals from the brain from unwanted signals, including environmental fields and artefacts due to head movement (Brookes et al., 2021). Given the degree of head movement required to play the violin, this was considered important. 3) Triaxial sensing also ensures as uniform coverage of the brain as possible (Boto et al., 2022). Finally, our system was able to integrate multiple signals including MEG, motion tracking (of multiple independent bodies including the violin, bow and the participants head) and the sound of the violin, with the latter two being used to control the paradigm and provide accurate time windows for analysis. This integration was important for the present study and will likely become even more important in future naturalistic neuroscience experiments where monitoring what a participant is doing and time-locking it to the MEG data will become critical.

There are some limitations of the present system which should not be ignored. The biggest limitation of the system is the lack of field control. Although closed-loop sensor operation kept the sensors working and their outputs linear, head movements in the background field still cause significant artefacts that can obfuscate the MEG data. The use of large electromagnetic coils to reduce background fields across the entire head volume is well known to reduce this effect, significantly improving the quality of OPM-MEG data recorded with a wearable system (Rea et al., 2021). Here, at the time of recording, such field control was unavailable (due to the fact that the motion capture system was already controlling the paradigm in real time, and additional field control would have been challenging from a hardware point of view) . This was deemed acceptable since beta oscillations are relatively high frequency (in healthy adults) and motion artefacts typically manifest at lower frequencies. However, future studies of naturalistic paradigms should seek to use field control. A second limitation relates to sensor calibration; recent work has shown that the use of known magnetic fields to calibrate an OPM-MEG system prior to use can lead to improvements in source reconstruction (Hill et al., 2025; Schofield et al., 2025). Such techniques were unavailable at the time of the present data recordings, and whilst this is unlikely to have had a major effect on the results presented, it is likely that these results could have been improved by calibration techniques. Again, these techniques should be used in future studies.

Finally, we acknowledge some limitations of the task design. First, some aspects of our paradigm were uncontrolled – for example some participants could have felt muscle fatigue during the task which may have affected the second scanning session more than the first. It is therefore not impossible that beta effects could have been influenced by this. Second, the study included just a single violin lesson as “training”. This is the minimum possible intervention, and future OPM-MEG studies of this type should arguably seek to employ more expansive training programmes – for example spread out over several lessons or at least providing participants a chance to practice what they have learned (e.g. taking a violin home overnight). Nevertheless, the fact that subtle changes in beta oscillations were measurable with just a single lesson is compelling and speaks to the sensitivity of OPM-MEG as a means to investigate motor learning. Third, the way in which we measured “success” was subjective – by asking each participant to rate how well they played the tune using a 5-point Likert scale. In principle, it should be possible to use recorded sound to derive an objective measure of success. However, this is complex since one can measure the frequency of the notes, the clarity of the sound, the duration of notes, the overall time taken to play the tune and even the amount of musical expression etc. All of these factors might combine to a single metric of “success”. However, how they combine likely differs (subjectively) across participants. For this reason, we chose to use the simple self-assessment metric. Nevertheless, future studies should consider objective analyses of playing success.

## CONCLUSION

We have shown that OPM-MEG data can be recorded as individuals carry out a naturalistic motor learning task. As expected, playing the violin resulted in periods of movement, during which beta oscillations reduced in amplitude. This was followed by an increase in beta amplitude on movement cessation, again as expected. Moreover, when recording OPM-MEG data before and after a violin lesson, beta power whilst playing was found to be elevated after the lesson compared to before; the change in beta amplitude elicited by playing was also more pronounced after the lesson. These findings fit well with current models of beta dynamics, suggesting that beta amplitude tends to be higher when an individual has more certainty of the movements that they are attempting to carry out, compared to when those movements are less well rehearsed. Our study adds to an expanding literature on the role of beta oscillations in brain function, and this is important given multiple findings of abnormal beta dynamics in disorders. The study also showcases the utility of OPM-MEG for neuroscientific research into naturalistic motor learning.

## ACKNOWLEDGEMENTS

This work was supported by an Engineering and Physical Sciences Research Council (EPSRC) Grant (EP/Z535722/1) and the UK Quantum Technology Hub in Sensors Imaging and Timing (QuSIT), funded by EPSRC (EP/Z533166/1). We acknowledge a Medical Research Council (MRC) Mid-Range Equipment grant (MC_PC_MR/X012263/1).

## DATA AVAILABILITY STATEMENT

Data and code will be made freely available upon acceptance of this manuscript, via Zenodo.

## SUPPLEMENTARY INFORMATION

**Figure S1:**
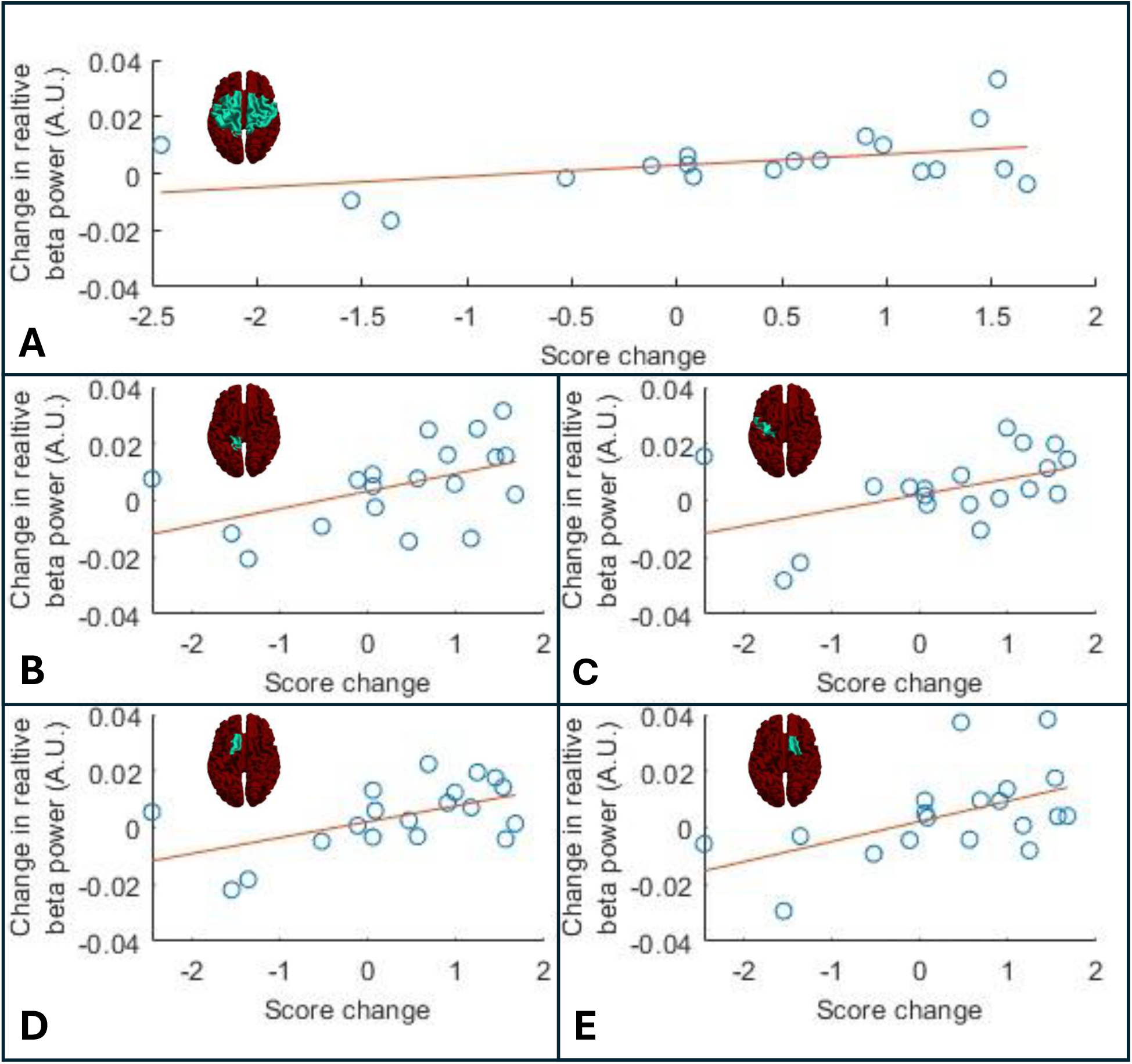
Correlations between the change in relative beta power during the playing window between sessions, and the change in trial averaged self-assessment score. A) Beta power in the whole motor network (r = 0.3; p = 0.071). B) left dorsolateral motor cortex (r = 0.47; p = 0.04). C) left dorsomedial somatosensory cortex (r = 0.34; p = 0.034); D) Left dorsomedial premotor area (r = 0.55; p = 0.014). E) Right dorsomedial premotor area (r = 0.52; p = 0.02).

## APPENDIX 1 Examples of bad trial rejection

**Figure A1:**
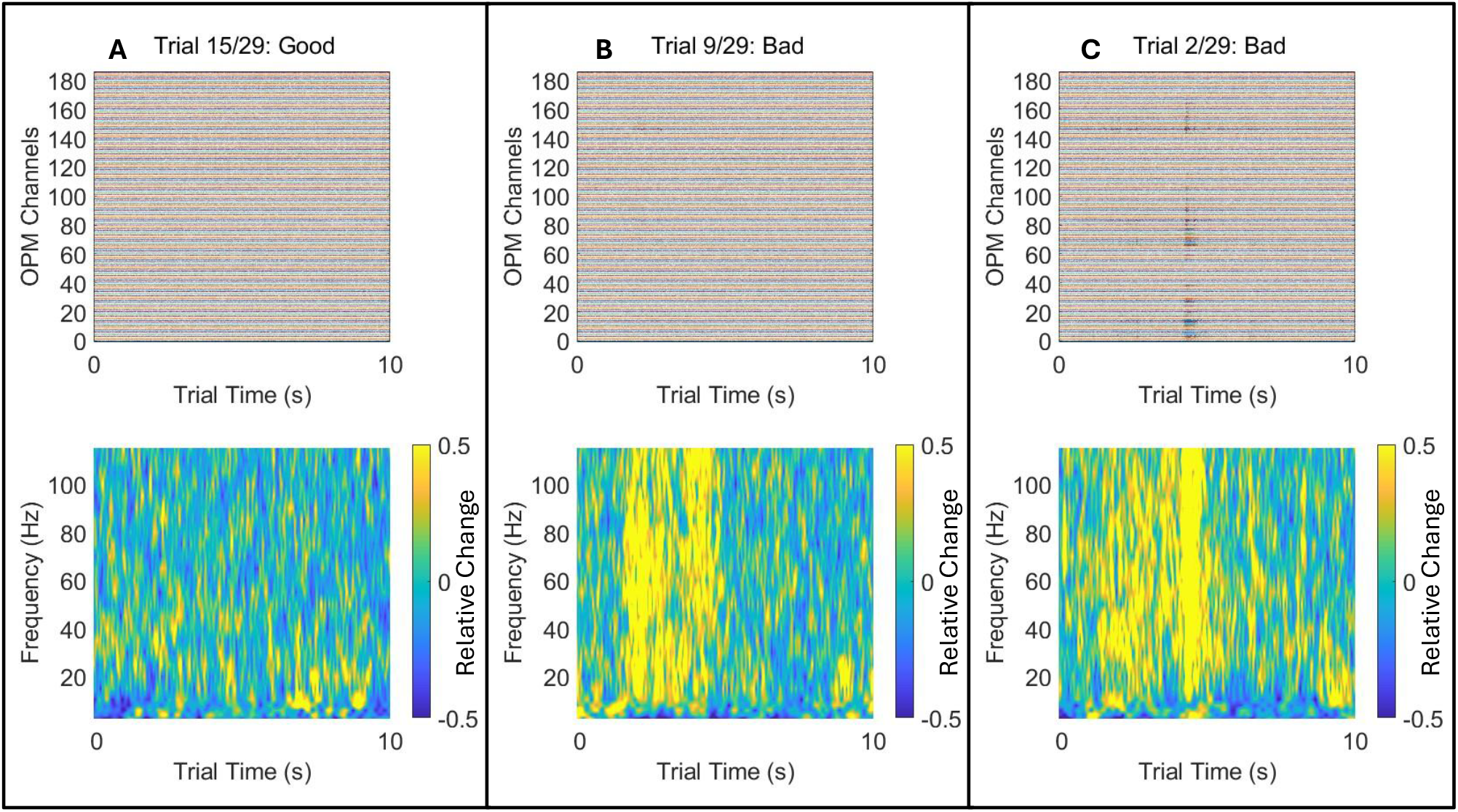
Examples of bad trial rejection. Upper panels show trial data for all MEG channels (filtered 1-150 Hz and following tSSS); lower panel shows time frequency data (averaged across channels). A) A good trial; B) A bad trial where the channel data look fine but muscle artefact is clear in the time frequency data; C) A bad trial where both the time frequency and channel data exhibit interference. These data are shown for a representative 10 s window.

A challenge for OPM-MEG applied to naturalistic experimental paradigms is how to deal with artefacts. In particular, when movement is encouraged, electrical activity in muscles (especially in the face, neck and shoulders) generates interference in MEG data which can obfuscate real brain activity. Here we chose to deal with this by removing ‘bad’ trials. Visual data inspection was carried out in two ways: 1) by examining the MEG data for all channels and 2) by combining time frequency data across channels. Figure A1 shows three examples of single trial data. In all three examples, the channel data (filtered 1-150 Hz with tSSS applied) are shown in the upper panel, whilst the time frequency data (averaged across channels) are shown in the lower panel. Panel A shows an example of a good trial; panel B shows an example of a trial where the channel data do not show obvious interference but muscle artefact is clear in the time frequency data; panel C shows data where interference is clear in the time-frequency and channel data. The trials shown in B and C were removed from the final analysis.

## APPENDIX 2 Regional beta power differences within the sensorimotor network

Data in Figure 3 shows a significant increase in beta power in the sensorimotor network, after the lesson compared to before. This was the case for both the playing and rest windows. However, in our primary analysis, we had collapsed data across all regions of the sensorimotor network (to avoid a multiple comparisons problem). As a post-hoc analysis, we also assessed how this significant change varied across the 12 different MarsAtlas regions within the sensorimotor network.

Figure A2 shows results of this analysis. The upper panel shows the playing window whilst the lower panel shows rest window. In both cases the height of the bars show the mean (across participants) beta change (post lesson minus pre-lesson) for the 12 regions that formed our sensorimotor network (these data are also shown as a colour plot, by the inset images in Figure 3 (right hand panel)). Error bars show standard deviation over participants and the individual data points show each participant. Inset brains show the locations of each of the 12 regions (which are also labelled on the graph).

**Figure A2:**
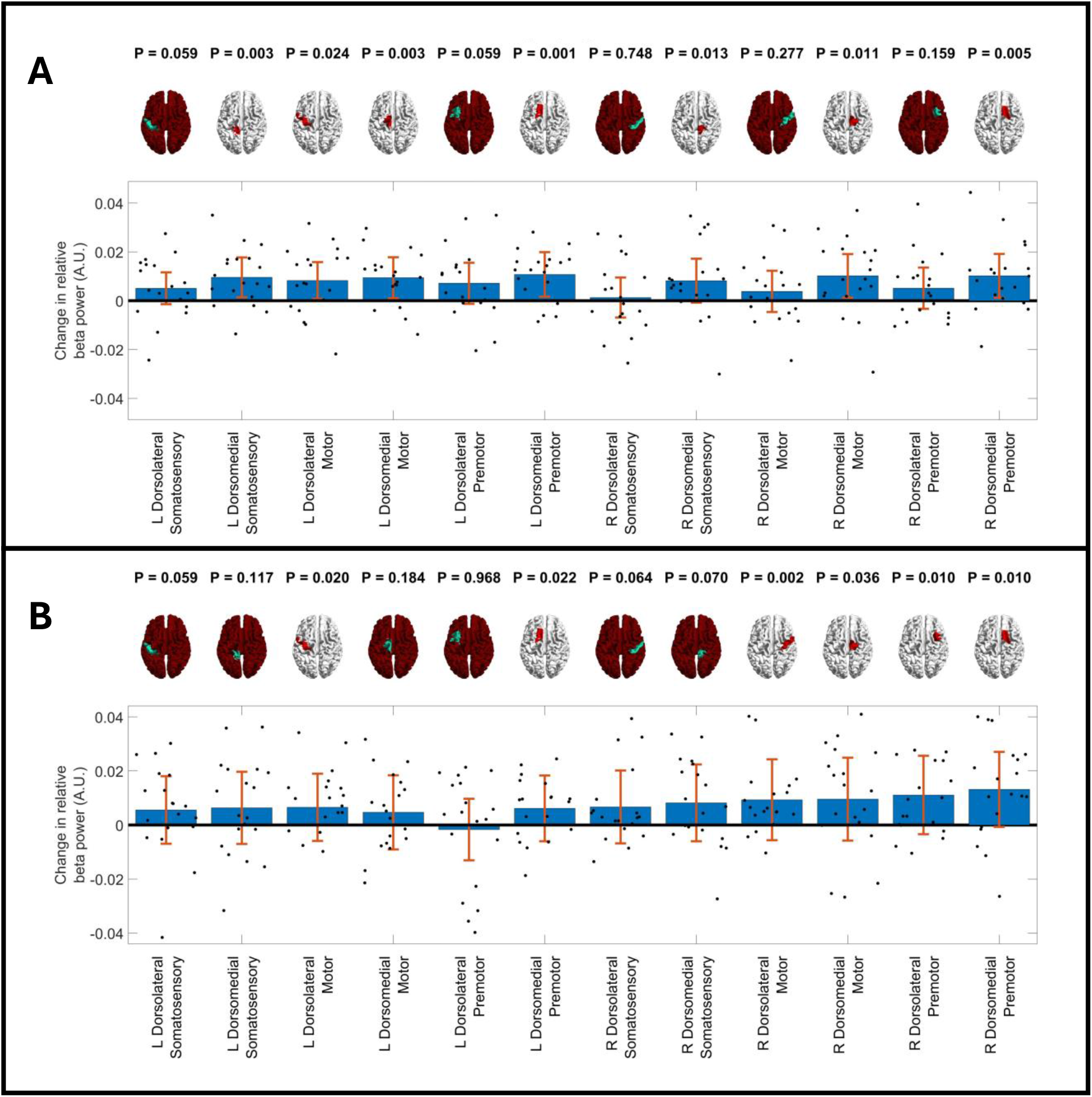
Beta band change before and after the lesson, divided by brain region. A) Playing Window. B) Rest window. In both cases the bar chart shows mean (across participants) beta change across 12 regions in the sensorimotor network. Error bars show standard deviation over participants and the individual data points show data for each participant. Inset brains show the locations of each region. P-values represent independent statistical tests for each region.

In agreement with Figure 3, most regions show a positive change between the two sessions, indicating the general trend for larger beta power after the lesson. During the playing window, this effect was largest in the left and right dorsomedial premotor cortices, areas that are associated with motor planning. Similarly large effects were also seen in left and right dorsomedial motor cortex. Interestingly, both left and right dorsolateral somatosensory cortex showed no measurable change in beta power. Similarly the left and right dorsolateral premotor and right dorsolateral motor cortex showed no significant change.

During the rest window, change in beta power was strongest in the right hemisphere including dorsomedial premotor, dorsolateral premotor, dorsomedial motor and dorsolateral motor cortices. Only the dorsomedial premotor cortices and dorsolateral motor cortices showed a significant effect in left hemisphere. This right hemispheric dominance may relate to the fact that the left hand is most heavily engaged in the task (i.e. moving the fingers over the strings).

In the main paper, we chose to run the statistical test over the whole motor network (rather than dividing the network into its 12 constituent regions) to avoid a multiple comparison issue. It is however noteworthy that had we run a statistical test in all 12 regions, and corrected for multiple comparisons using the Benjamini Hochberg method, then during the playing window, 7 of the 12 regions would remain showing significant beta increase after the lesson, even following multiple comparison correction. In the rest window, 2 of the 12 regions would remain following the same multiple comparison correction.

